# *Drosophila* FMRP recruits the miRISC to target mRNAs to repress translation

**DOI:** 10.1101/2023.05.03.539280

**Authors:** Navneeta Kaul, Sarala J. Pradhan, Nathan G. Boin, Madeleine M. Mason, Julian Rosales, Emily L. Starke, Emily C. Wilkinson, Erich G. Chapman, Scott A. Barbee

## Abstract

Fragile X syndrome (FXS) is the most common inherited form of intellectual disability and is caused by mutations in the gene encoding for the Fragile X messenger ribonucleoprotein (FMRP). FMRP is an evolutionarily conserved and neuronally enriched RNA binding protein (RBP) with functions in the control of processes including RNA editing, RNA transport, and protein translation. Specific target RNAs play critical roles in neurodevelopment including the regulation of neurite morphogenesis, synaptic plasticity, and cognitive function. The different biological functions of FMRP are modulated by its cooperative interaction with distinct sets of neuronal RNA and protein binding partners. Here, we focus on interactions between FMRP and components of the microRNA (miRNA) pathway. Using the *Drosophila* model system, we show that dFMRP can repress the translation of a reporter mRNA via a deadenylation-independent mechanism. This repression requires the activity of both AGO1 and GW182, conserved components of the miRNA-containing RISC (miRISC). Interestingly, we find that dFMRP can bind directly to a short stem loop structure in the reporter and that dFMRP binding is a prerequisite for repression by miR-958. Finally, we show that *dFmr1* interacts genetically with *GW182* to control neurite morphogenesis. Collectively, these data suggest the dFMRP can directly recruit the miRISC to nearby miRNA binding sites and then repress translation via the activity of the miRISC effector, GW182.

## INTRODUCTION

Fragile X Syndrome (FXS) is the most common inherited neurodevelopmental disorder in humans and is the leading monogenetic cause of autism (Santoro et al. 2012). Most cases of FXS are caused by expansion of the CGG trinucleotide repeat (>200) in the 5’ untranslated region (UTR) of the X-linked *FMR1* gene, which leads to DNA hypermethylation and epigenetic transcriptional silencing. *FMR1* encodes for the Fragile X messenger ribonucleoprotein (FMRP), an evolutionarily conserved RNA binding protein (RBP) that has been implicated in multiple steps of RNA metabolism including direct roles in RNA editing, translation, and RNA transport (Lai et al. 2020; Richter and Zhao 2021). FMRP contains three canonical RNA binding motifs including two heterogenous nuclear ribonucleoprotein K homology (KH) domains and an arginine-glycine-glycine (RGG) motif (Richter and Zhao 2021). In numerous studies, FMRP has been shown to bind to 1000s of mRNAs in the brain, many of which are involved in the regulation of important processes such as neurite morphogenesis and synaptic plasticity (Chen and Joseph 2015).

FMRP has been best characterized as a repressor of translation and the loss of translational control is widely believed to be directly linked to the deficits seen in FXS (Darnell and Klann 2013). FMRP has been shown to repress translation via several distinct mechanisms. First, mammalian FMRP can repress cap-dependent translation in specific target mRNAs by interacting with the Cytoplasmic FMRP Interacting Protein (CYFIP) (De Rubeis et al. 2013). Second, FMRP has been best characterized as a repressor of translational elongation. FMRP co-sediments with polyribosomes in sucrose gradients suggesting that it may cause the reversible stalling of ribosomes on target mRNAs (Feng et al. 1997). In humans, FMRP can inhibit elongation *in vitro* via its RGG motif and disordered C-terminal domain (Athar and Joseph 2020). In contrast, cryo-EM studies in *Drosophila* have revealed that the KH1 and KH2 domains can directly interact with the L5 protein on the 80S ribosome to block translation elongation (Chen et al. 2014).

FMRP has also been found to repress the translation of specific target mRNAs by interacting with conserved components of the microRNA (miRNA) pathway (Kenny and Ceman 2016). First, mammalian FMRP can reversibly interact with the riboendonuclease, Dicer, and is thought to modulate the processing of precursor miRNAs (pre-miRNAs) into mature miRNAs (Cheever and Ceman 2009). Second, mammalian and *Drosophila* FMRP interact biochemically with the Argonaute (AGO) proteins, core components of the RNA-induced silencing complex (RISC) (Jin et al. 2004; Lee et al. 2010). The AGO proteins function to bind to small RNAs (such as miRNAs) and facilitate their interaction with specific sequences in target mRNAs. Finally, mammalian FMRP binds to the RISC-associated RNA helicase, MOV10 (Kenny et al. 2014). Interestingly, the FMRP/MOV10 complex has been shown to both block and activate the translation of distinct target mRNAs by regulating the ability of the miRNA-containing RISC (miRISC) to bind to nearby target sequences (Kenny and Ceman 2016; Kenny et al. 2020).

A critical outstanding question that remains in the field is understanding how FMRP represses translation of bound mRNAs by recruiting components of the miRNA pathway. The AGO proteins are alone insufficient to mediate the miRNA-mediated silencing of target mRNAs (Niaz and Hussain 2018). Instead, silencing is facilitated by the AGO-associated GW182 proteins. GW182 is effector in the miRISC and acts as a scaffold to recruit proteins required for translational repression or mRNA deadenylation followed by 5’-to-3’ exonucleolytic decay (Niaz and Hussain 2018). In this study, we have developed a novel functional assay to better understand how the *Drosophila* ortholog of FMRP (dFMRP) represses translation via the miRNA pathway. We found that tethered dFMRP can repress reporter translation through a process that does not involve ribosome-scanning or deadenylation. Interestingly, repression of the reporter is abrogated by the depletion of both AGO1 and GW182 by RNA interference (RNAi). Conversely, repression of the reporter by exogenously expressed miR-958 first requires FMRP binding. Finally, we show that *dFmr1* interacts genetically with *GW182* to regulate synapse morphogenesis *in vivo*. Taken together, our study suggests that an FMRP/AGO/GW182 complex may target critical neuronal mRNAs for translational repression via a deadenylation- and decay-independent mechanism.

## RESULTS

### dFMRP can repress reporter translation in a tethered functional assay

To better understand how dFMRP regulates translation, we turned to a functional tethering assay we recently adapted to study dFMRP activity in *Drosophila* Schneider (S2) cells (Starke et al. 2022). Full length dFMRP was fused to a N-terminal λN-HA tag and co-expressed with a firefly luciferase (FLuc) reporter containing five tandem λN binding sites (BoxB hairpins) (Fig. 1A). These were co-expressed with a *Renilla* luciferase (RLuc) reporter as a transfection control. S2 cells were transfected with increasing concentrations of tagged dFMRP or a λN-HA control (Fig. 1B). As expected, expression of full length dFMRP significantly repressed reporter expression at all concentrations tested (Fig. 1C) (Starke et al. 2022). This repression was at the level of translation because the amount of polyadenylated mRNA did not similarly decrease (Fig. 1C). To further understand mechanistically how dFMRP repressed translation, we next assessed the translation status of the reporter mRNA using polysome profiling in cells transfected with 0.5 μg of the plasmid expressing dFMRP or the control (Supplemental Fig. S1). Interestingly, we found that there was a notable shift in the enrichment of reporter mRNA to light polysome fractions after expression of dFMRP (Fig. 1D). This shift is characteristic of mRNAs that are poorly translated or whose translation has been disrupted at the elongation step (Chasse et al. 2017).

**Figure 1.**
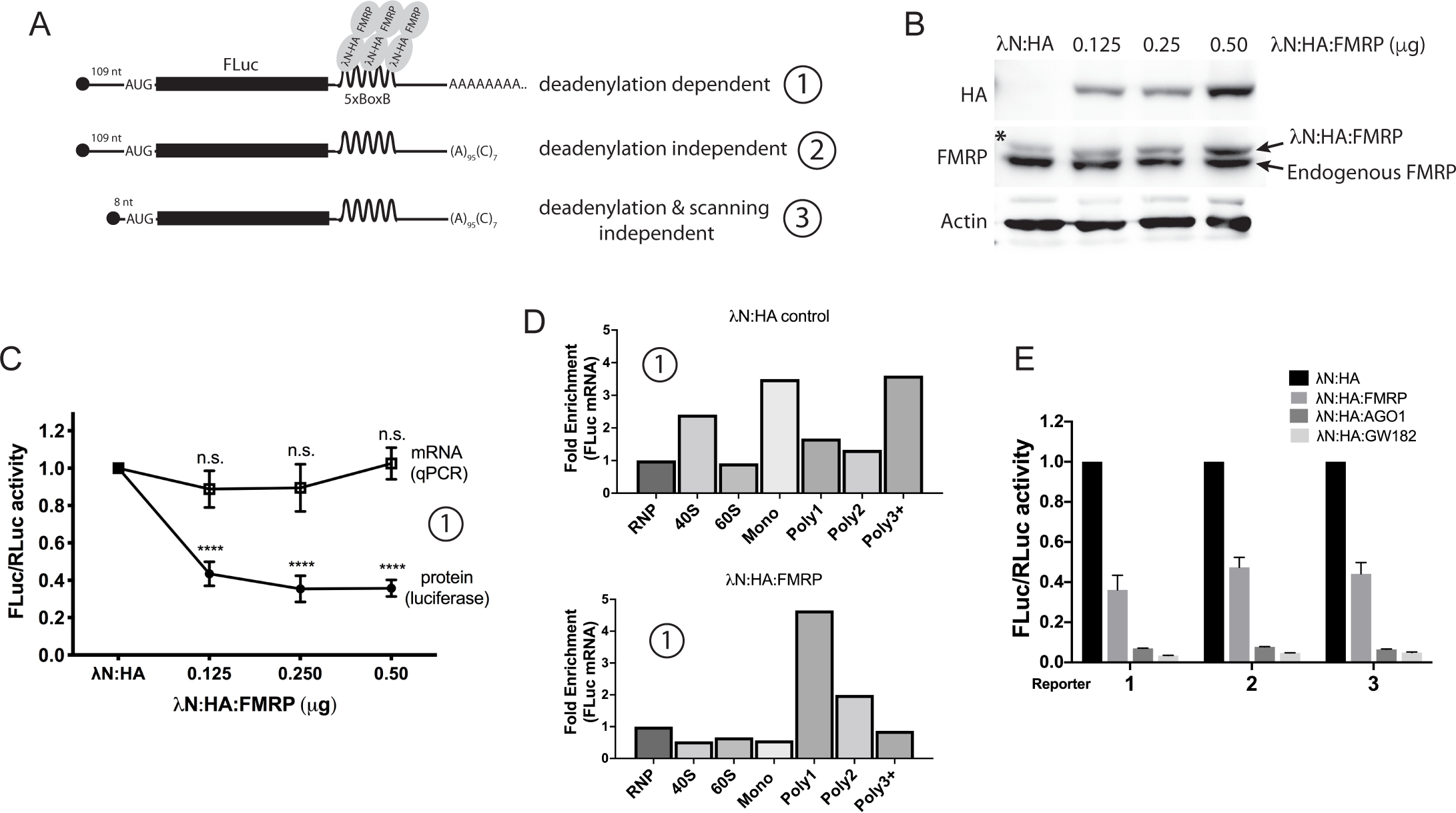
FMRP silences translation of reporter mRNAs via a deadenylation- and scanning-independent mechanism. (A) Schematic representation of the translational reporters used in Figure 1 and 2. (B and C) Tethering assay in S2 cells transfected with FLuc-5xBoxB reporter #1 and increasing concentrations of a plasmid expressing λN-HA tagged dFMRP. (B) Western blot showing levels of λN-HA-dFMRP relative to endogenous FMRP and an Actin loading control. (C) FLuc activity and RNA levels were normalized to those of the RLuc transfection control and then shown relative to the λN-HA peptide negative control after its values were set to 100%. The asterisk marks a band in the control corresponding to a heavier isoform of endogenous dFMRP. (D) FLuc reporter #1 RNA abundance in fractions collected for polysome profiling. Relative fold enrichment was calculated relative to RNA levels in the small RNP fraction for each condition. (E) FLuc activity of the indicated reporters after tethering FMRP, AGO1, and GW182. Results for each reporter were normalized and described for (C). All luciferase assay and qPCR experiments were done in three biological replicates. Statistical significance for results in (C) was determined by one-way ANOVA followed by Dunnett’s multiple comparison test. **** p < 0.0001.

Finally, we analyzed the ability of dFMRP to repress translation using two additional reporters (Fig. 1A). The first contained a 3’ end generated by a self-cleaving hammerhead ribozyme (HhR) that lacks a poly(A) tail and consequently cannot be targeted for deadenylation (Zekri et al. 2013). The second lacked a poly(A) tail but also contained a short 5’ UTR (8 nucleotides) which can initiate translation without ribosome scanning (Kuzuoglu-Ozturk et al. 2016). Interestingly, full length dFMRP was capable of similarly repressing all reporters, reducing translation to ∼ 40% of λN-HA controls (Fig. 1E). Taken together, these data suggest that tethered dFMRP can repress reporter translation via a ribosome scanning- and deadenylation-independent mechanism. Collectively, these data provide support for a model where dFMRP is blocking translation of the reporter mRNA, presumably at the level of translational elongation.

### dFMRP represses reporter translation via conserved components of the miRNA pathway

The *Drosophila* miRNA proteins AGO1 and GW182 have been shown to regulate all FLuc reporters in a similar manner to what we show for dFMRP (Kuzuoglu-Ozturk et al. 2016). To verify this, we obtained and tested plasmids expressing λN-HA tagged full length AGO1 and GW182 in the tethering assay. As expected, we found that both AGO1 and GW182 significantly repressed translation of the reporter and that results were more robust than those for dFMRP (Fig. 1D). Because an evolutionarily conserved genetic and biochemical interaction between FMRP and the miRNA pathway is well established (Zhou et al. 2019), we next asked whether repression of the reporter by dFMRP required components of the miRNA pathway. We initially focused on the role of GW182 because it is the effector in the miRISC and directly induces translational repression while AGO1 is insufficient for silencing (Niaz and Hussain 2018). To examine this interaction, we co-transfected S2 cells with λN-HA tagged dFMRP, FLuc and RLuc reporters, and double stranded RNA (dsRNA) targeting two regions of the *GW182* transcript. Surprisingly, we found that knockdown of *GW182* expression with both dsRNAs led to significant derepression of reporter translation by dFMRP (Fig. 2A-B). To further explore this interaction, we expanded our analysis to include additional conserved components of the miRNA pathway that have been shown to interact with FMRP in both *Drosophila* and mammals. As with *GW182*, we found that knockdown of *Ago1* expression resulted in significant de-repression of reporter translation (Fig. 2C-D). In contrast, knockdown of *Dcr1* expression trended towards, but surprisingly did not reach, statistical significance (Fig. 2C-D). It is possible that knock down of *Dcr1* may not sufficiently deplete pre-existing miRISC complexes. Lastly, co-transfection with dsRNA targeting GFP (not expressed in this system) had no impact on translational repression by FMRP (Fig. 2C). Collectively, these data suggest that reporter repression by FMRP requires components of the miRISC.

**Figure 2.**
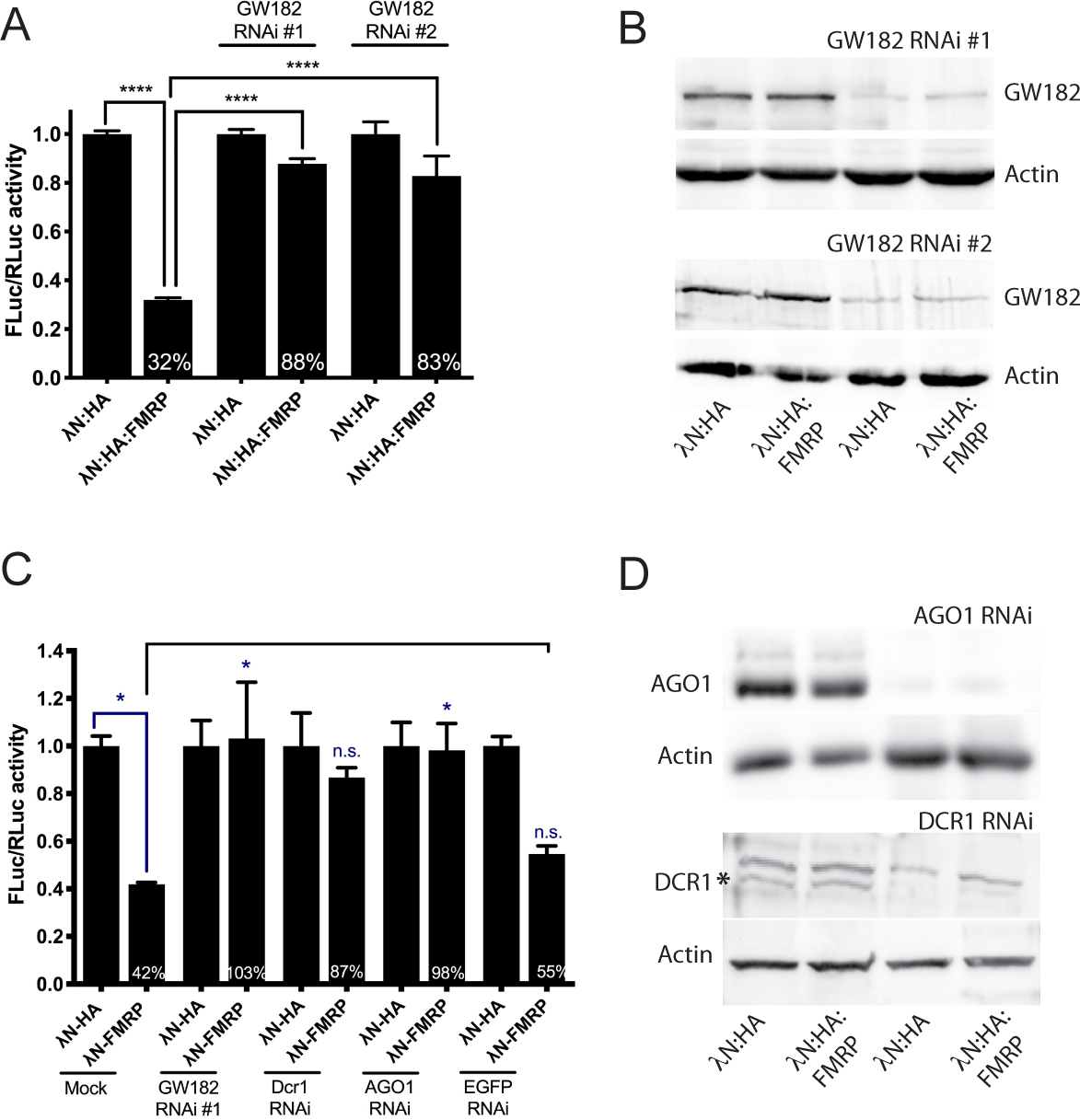
Reporter mRNA repression by tethered FMRP requires conserved components of the miRNA pathway. (A) FLuc activity in S2 cells transfected FLuc-5xBoxB reporter #1, λN-HA tagged dFMRP, and dsRNA targeting two non-overlapping regions of the *GW182* transcript. (B) Western blot showing levels of GW182 expression with and without co-transfected dsRNA. (C) FLuc activity in S2 cells transfected FLuc-5xBoxB reporter #1, λN-HA tagged dFMRP, and dsRNA targeting the mRNAs encoding for GW182, DCR1, AGO1, and EGFP (negative control). D) Western blot showing levels of AGO1 and DCR1 expression with and without co-transfected dsRNA. The asterisk marks the 255 kDa DCR1 protein. As there are no other predicted isoforms of DCR1 in *Drosophila*, and it is not affected by *Dcr1* RNAi, the higher band is likely nonspecific. All experiments shown were done in triplicate. Statistical significance for results in A and C was determined by one-way ANOVA followed by a Tukey’s multiple comparison test. Significance between specific conditions is indicated by brackets. * p < 0.05, **** p < 0.0001.

### AGO1 colocalize strongly with FMRP-containing cytoplasmic granules

It is well described that FMRP associates with membraneless cytoplasmic granules that contain RBPs and translationally repressed mRNAs (Lai et al. 2020). Based on the requirement for the miRNA pathway to repress reporter translation, we hypothesized that, as seen with human orthologs, miRISC proteins may colocalize with dFMRP in these granules (Lee et al. 2010). To test this, we co-transfected S2 cells with a plasmid that expressed a GFP-tagged full length dFMRP protein along with one that expressed mCherry-tagged miRNA pathway proteins (Fig. 3A). As expected, dFMRP colocalized very strongly with AGO1 in S2 cells (Fig. 3A-B). In contrast, while most dFMRP granules do not contain GW182 and Dcr1, a significant population of AGO1- and GW182-containing granules colocalized with dFMRP (Fig. 3C). Finally, it has been shown that dFMRP interacts biochemically with AGO1 although its association with GW182 has not been investigated (Jin et al. 2004). Moreover, the dependence of this association on RNA is not known. To address these questions, we co-transfected S2 cells with plasmids expressing HA-tagged dFMRP and either FLAG-tagged AGO1 or GW182. Consistent with published results, AGO1 co-immunoprecipitated strongly with dFMRP (Jin et al. 2004; Lee et al. 2010) but we found that this interaction is not dependent upon RNA (Fig. 3D). Similarly, we show that GW182 also interacted with dFMRP in an RNA-independent manner (Fig. 3D). This result is not unexpected as human GW182 binds directly to AGOs within the miRISC, and GW182 is believed to act as a bridge between AGOs and downstream effectors (Jonas and Izaurralde 2015; Elkayam et al. 2017). Together, these data suggest a protein-protein interaction between dFMRP and the miRISC.

**Figure 3.**
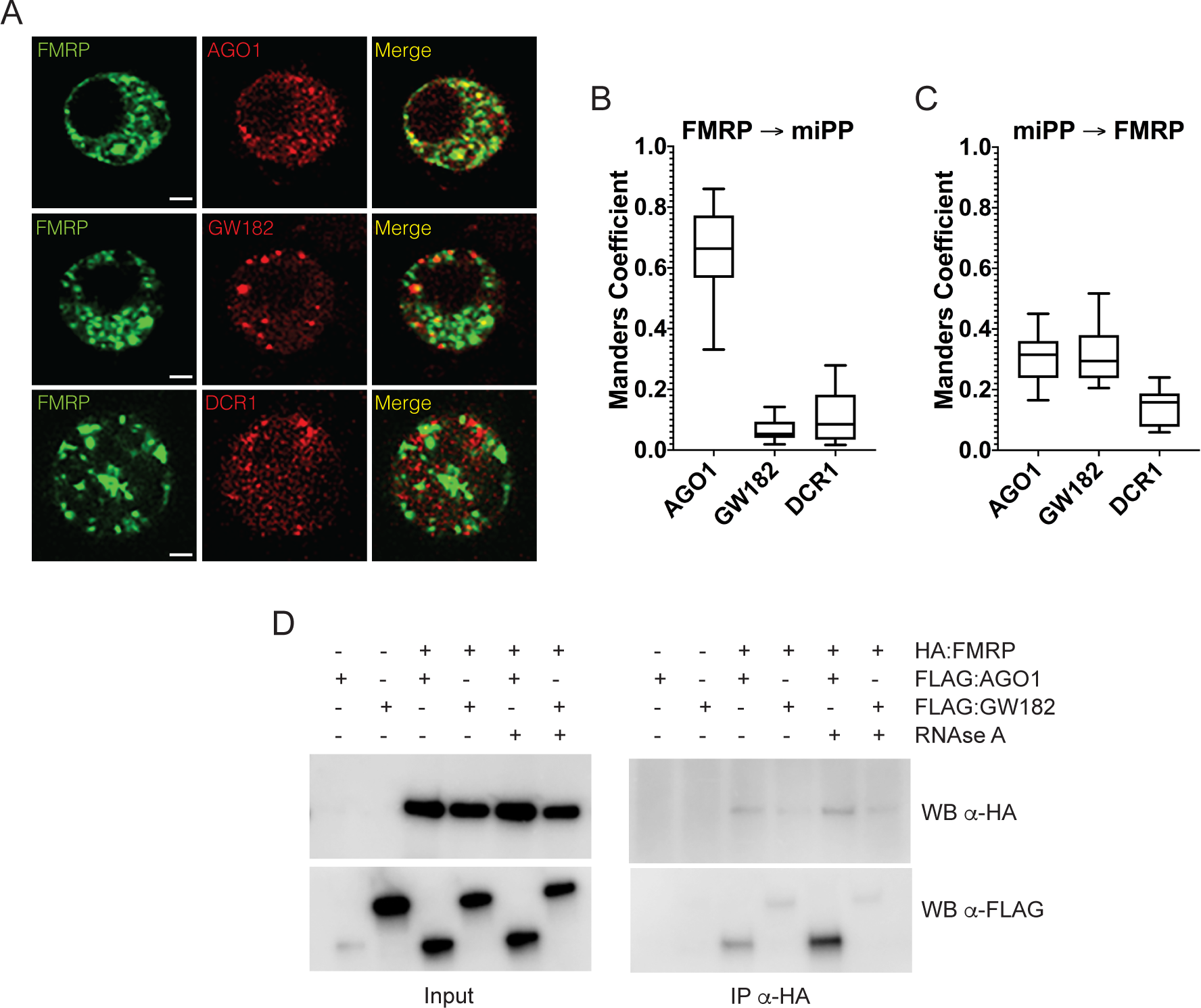
FMRP colocalizes and biochemically interacts with component of the miRISC. (A) Confocal micrographs of representative S2 cells transfected with EGFP-tagged dFMRP (green) and the indicated mCherry-tagged miRNA pathway protein (miPP; red). Merged images are included for visual comparison of the colocalization (or lack of colocalization) between proteins. Scale bars = 2 μm. (B and C) Graphs showing the degree of overlap as determined by Manders correlation coefficients between green FMRP pixels that also contain red miPP fluorescence (B) and *vise versa* (C). Number of cells analyzed for colocalization were: FMRP/AGO1 (n = 12), FMRP/GW182 (n = 14), and FMRP/DCR1 (n = 13). (D) Western blots showing results of co-immunoprecipitaton assays in S2 cells cotransfected with λN-HA tagged dFMRP and FLAG-tagged AGO1 or GW182. To determine if RNA is required to facilitate interactions between proteins, some reactions were incubated with RNAse A during the immunoprecipitation step.

### dFMRP can bind directly to the BoxB hairpin sequence and repress reporter translation

Mouse and human FMRP has been shown to bind several sequence and structural elements in target transcripts via its RNA binding domains. These motifs include G-quadraplexes via its RGG box domain (Ramos et al. 2003). Additionally, the KH domains have been shown to interact with a RNA pseudoknot structure and two short sequences that are enriched in FMRP bound transcripts, CGGA and ACUK (where K = G/U and W = A/U) (Darnell et al. 2005; Ascano et al. 2012). Interestingly, FMRP has also been shown to bind to a structural element consisting of three tandem stem loops in the *Sod1* mRNA and regulate its translation (Bechara et al. 2009). Based on this, we hypothesized that dFMRP may be able to bind directly to the BoxB stem loop within the 3’UTR of the reporter transcript. To test this, we first conducted an electrophoretic mobility shift assay (EMSA) using full length dFMRP protein and a minimal RNA probe consisting of a single BoxB stem loop (Fig. 4A). Surprisingly, we observed a clear, concentration-dependent shift in the mobility of the labeled probe (Fig. 4B). On the other hand, dFMRP did not cause a shift in the presence of excess concentrations of an unstructured control RNA (Fig. 4B). These data confirmed that untethered FMRP could bind specifically to the BoxB stem loop.

**Figure 4.**
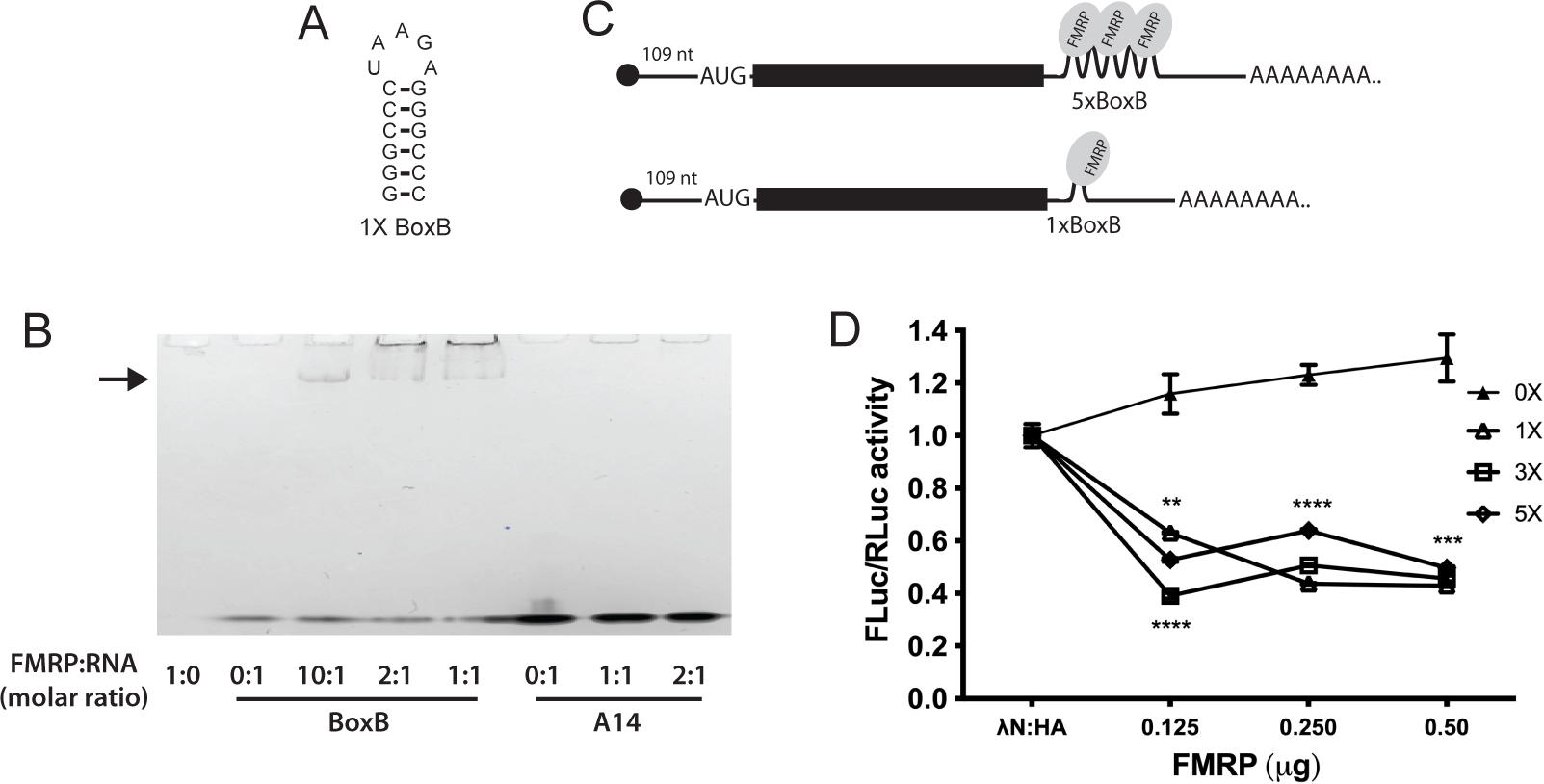
Untagged and untethered FMRP can bind directly to the BoxB stem loop to repress reporter translation. (A) The predicted structure of the 1xBoxB stem loop. (B) A “cold” native gel showing results of the EMSA experiments. Purified dFMRP was incubated with *in vitro* transcribed RNAs corresponding to the minimal 1xBoxB stem loop or an A14 unstructured control at the indicated molar ratio. The bands and the bottom of the gel show the unbound RNA probes. The arrow indicates supershifted 1xBoxB probe bound to dFMRP. (C) Schematic representation of two translational reporters used in Figure 4D. (D) Luciferase assay in S2 cells transfected with the indicated FLuc-BoxB reporters and increasing concentrations of a plasmid expressing untagged dFMRP. Statistical significance was determined by two-way ANOVA followed by Dunnett’s multiple comparison test. ** p < 0.01, *** p < 0.001, **** p < 0.0001.

We next asked if dFMRP could repress the expression of the luciferase reporter containing five tandem BoxB stem loops (5xBoxB). We co-transfected S2 cells with the reporter and increasing concentrations of a plasmid expressing dFMRP lacking the λN-HA tag. We observed that untethered FMRP was able to significantly repress reporter expression, even at the lowest concentrations tested (Fig. 4D). To examine this interaction further, we constructed a series of reporters that contained three, one, or no tandem BoxB stem loops (3xBoxB, 1xBoxB, and 0xBoxB) and similarly co-transfected S2 cells with an increasing concentration of dFMRP (Fig. 4C). Interestingly, dFMRP was able to repress translation of the 1xBoxB reporter as efficiently and it did the 5xBoxB reporter (Fig. 4D). In comparison, removal of all BoxB stem loops abolished the ability of dFMRP to repress expression (Fig. 4D). Together, this suggests that endogenous dFMRP can repress translation of the reporter by binding directly to the BoxB stem loop.

### Repression of reporter translation by miR-958 requires dFMRP binding to the BoxB motif

The discovery that dFMRP could directly bind and repress the reporter allowed us to next determine whether repression of reporter expression via the miRNA pathway required dFMRP. We first used a bioinformatic approach to identify putative miRNA binding sites located near the within the SV40 3’UTR. Surprisingly, we found a strong predicted binding site for miR-958 located ∼ 60 nucleotides downstream of the 1xBoxB stem loop (Fig. 5A; seed = 7mer-m8, ΔΔG = −12.7). We have previously shown that miR-958 regulates activity dependent axon terminal growth at the *Drosophila* larval NMJ (Nesler et al. 2013). To address this question, we next asked whether the 1xBoxB reporter was a target for repression by miR-958. We co-transfected S2 cells with the 1xBoxB reporter and a plasmid expressing the primary miRNA (pri-miRNA) sequence encoding for miR-958 and found that reporter expression was significantly reduced (Fig. 5B). Then, to determine if repression by miR-958 required dFMRP activity, we co-transfected cells with the 1xBoxB reporter, miR-958, and dsRNA targeting the *dFmr1* transcript. Knockdown of *dFmr1* expression completely disrupted the ability of miR-958 to repress reporter expression (Fig. 5B-C). To confirm this result, we next asked if miR-958 was capable of repressing translation of the 0xBoxB reporter, which retains the predicted miR-958 binding site but is lacking the BoxB stem loop (Fig. 5A). As with *dFmr1* knockdown, removal of the BoxB stem loop abolished the repression activity of miR-958 (Fig. 5B). Collectively, these data strongly suggest that the translational repression activity of miR-958 first requires dFMRP binding to the BoxB stem loop.

**Figure 5.**
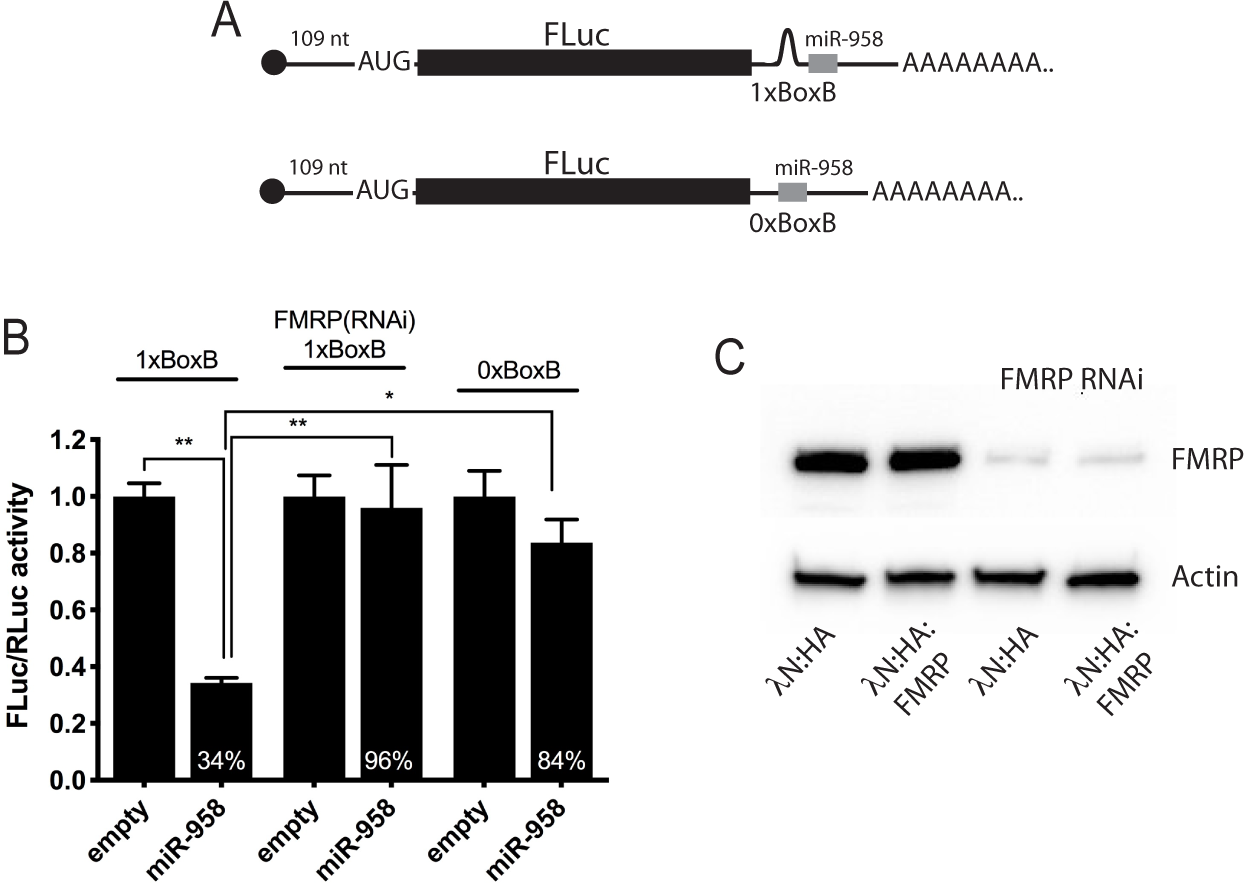
Reporter mRNA repression by miR-958 requires FMRP and the BoxB stem loop. A) Schematic representation of the translational reporters used in Figure 5 with the predicted binding site for miR-958. (B) FLuc activity in S2 cells transfected with the indicated FLuc-BoxB reporter (0xBoxB or 1xBoxB), a plasmid expressing the primary miR-958 (or an empty vector control), and (as indicated) dsRNA targeting the mRNA encoding for dFMRP. (C) Western blot showing levels of dFMRP expression with and without co-transfected dsRNA. Statistical significance was determined by one-way ANOVA followed by a Tukey’s multiple comparison test. Significance between specific conditions is indicated by brackets. * p < 0.05, ** p < 0.01.

### dFmr1 interacts genetically with GW182 to regulate NMJ morphogenesis

Loss of *dFmr1* expression causes strong defects in synapse structure and function at the *Drosophila* larval NMJ (Zhang et al. 2001). Furthermore, *dFmr1* interact genetically with *Ago1* to regulate synaptic growth and structure at the NMJ (Jin et al. 2004). Specifically, larvae heterozygous for both *dFmr1* and *Ago1* exhibit overgrowth and overelaboration of synaptic terminals, a phenotype like that seen following *dFmr1* loss-of-function (Jin et al. 2004). Both AGO1 and GW182 were co-expressed in motor neuron cell bodies in the larval CNS (Supplemental Fig. S2). Based on these observations, we sought to determine if *dFmr1* similarly interacted genetically with *GW182* to regulate NMJ morphogenesis. The *Drosophila* larval NMJ is used as a genetic model for the study of glutamatergic synapses in the mammalian brain (Schuster 2006). Foremost among its useful properties, its terminal synaptic boutons exhibit a high degree of plasticity during development (Collins and DiAntonio 2007). For this analysis, we examined the NMJs innervating muscle 6/7 in abdominal segment 3 of third instar larvae. This NMJ contains two types of glutamatergic boutons – 1b (big) and 1s (small) that are derived from two distinct neurons (Rohrbough et al. 2000). Type 1b boutons are highly plastic and can be distinguished by their larger size and higher levels of the postsynaptic density marker, Dlg (Lahey et al. 1994). To determine if *dFmr1* and *GW182* interact to regulate NMJ morphogenesis, we analyzed double heterozygotes (*dFmr1 −/+; GW182 −/+*) and compared results to single heterozygotes and a control for genetic background (Fig. 6). Synapse size was quantified by counting the number of type 1b synaptic boutons. Importantly, we observed a statistically significant increase in synaptic bouton number in *dFmr1 −/+; GW182 −/+* larvae relative to controls (Fig. 6F-G). This was seen despite the *GW182 −/+* single heterozygote having a strong overgrowth phenotype (Fig. 6D, G). As a positive control, we examined the genetic interaction between *dFmr1* and *Ago1*. Surprisingly, we observed a significant, but much weaker overgrowth phenotype then expected (Fig. 6E, G). However, differences between our results and those reported by Jin et al. (Jin et al. 2004) are likely due to differences in *dFmr1* alleles and normalization methods. Taken together, these data support a model where *dFmr1* requires components of the miRISC to regulate the normal development of glutamatergic synapses in the *Drosophila* model system.

**Figure 6.**
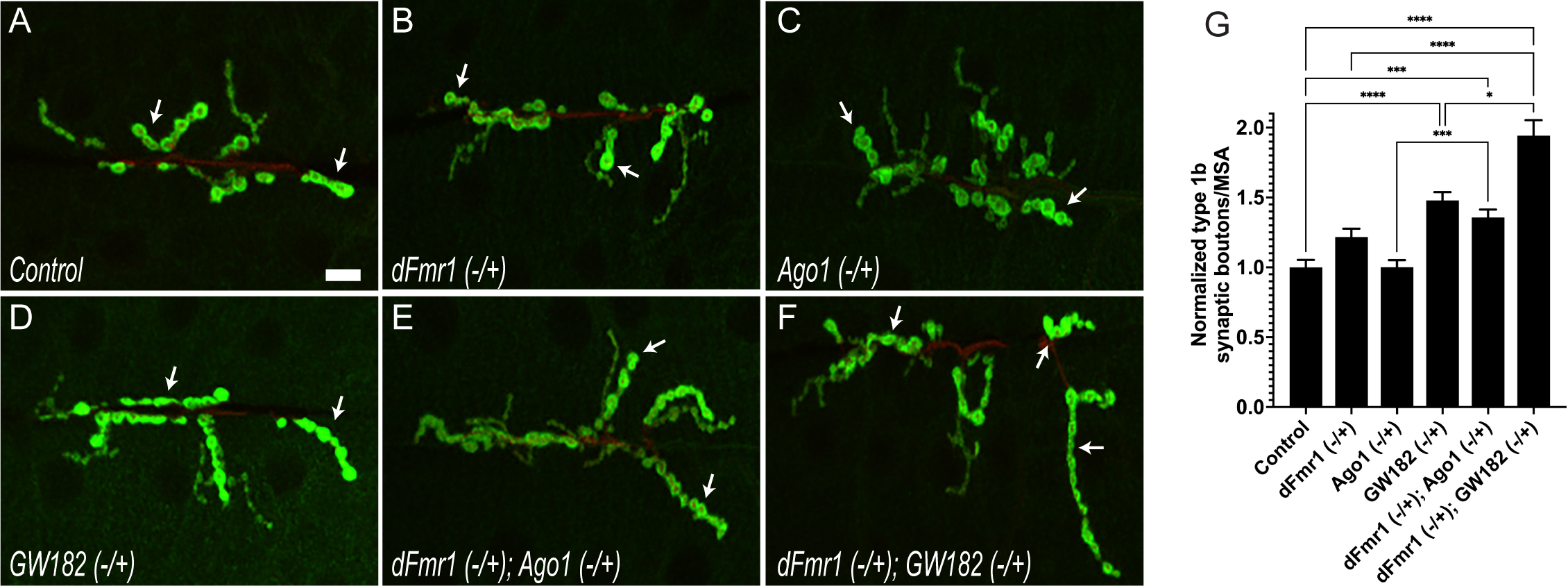
*dFmr1* interacts genetically with *GW182* to regulate neurite morphogenesis and the larval NMJ. (A-F) Representative images from the indicated genotypes of third instar larval NMJs immunostained with antibodies targeting DLG (green). Larger type 1b boutons are indicated by arrows. Scale bar = 10 μm. (G) The average number of type 1b synaptic boutons per NMJ for each genotype normalized to MSA. Values are shown relative to the *w^1118^ (Iso31)* negative control after its values were set to 100%. Statistical significance was determined using a Brown-Forsyth and Welch ANOVA followed by a Dunnett’s multiple comparison test. Significance between specific genotypes is indicated by brackets. * p < 0.05, *** p < 0.001, **** p < 0.0001.

## DISCUSSION

One critical function of FMRP is to regulate the translation of specific target mRNAs by blocking translational initiation, elongation, or acting through the miRNA pathway. However, what decides which of these mechanisms will be utilized by FMRP? Three lines of evidence suggest that this depends upon both combinatorial interactions with specific protein binding partners and/or sequence elements located within target mRNAs. First, FMRP can interact directly with CYFIP to regulate translational initiation (Napoli et al. 2008). This study shows that FMRP can recruit CYFIP to specific mRNAs, which in turn sequesters the cap-binding protein eIF4E and thereby preventing initiation in cap-dependent translation. However, it is not clear what specific sequence elements are bound by FMRP in target mRNAs to facilitate this process. Second, FMRP can interact directly with the ribosome to inhibit translation elongation. CLIP-seq experiments have shown that mammalian FMRP binds primarily to short sequence elements in coding sequence via its KH domains (Darnell et al. 2011). Evidence suggests that dFMRP may directly block ribosome translocation by inhibiting the interaction of tRNAs or elongation factors (Chen et al. 2014). Finally, a FMRP/MOV10 complex can bind to G-quadraplex sequences found in the 3’UTR of some target mRNAs. Interestingly, the FMRP/MOV10 interaction can protect a subset of bound mRNAs from AGO association, which increases levels of target mRNA expression (Kenny et al. 2020). In this case, the miRNA response element (MRE) is embedded within the G-quadraplex structure and MOV10 stabilizes the FMRP/quadraplex interaction.

Our data supports a simple model for translational repression by dFMRP via the miRNA pathway where FMRP/AGO1/GW182 complexes are directly recruited to a specific combination of sequence motifs located in the 3’UTR of target mRNAs (Fig. 7). These sequence motifs include both a binding site for FMRP and a nearby MRE, recognized by a specific miRNA. Importantly, the MRE is accessible and not embedded in G-quadraplex or complex secondary structure. GW182, the miRISC effector, then facilitates translation repression of the target mRNA.

**Figure 7.**
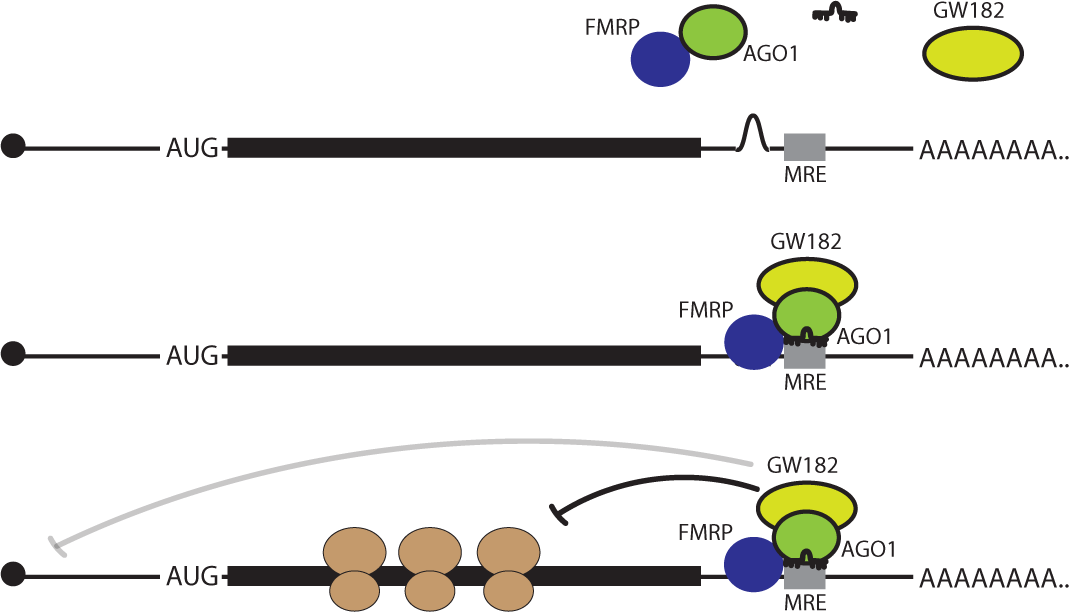
Model for repression of reporter mRNA expression by FMRP and the miRISC. Published and our data suggest that dFMRP may associate with unloaded AGO1 in cytoplasmic granules. After miRNA loading, the FMRP/AGO1/GW182 complex can be recruited to target mRNAs that have a FMRP binding site and nearby MRE. Once bound, GW182 facilitates the translational repression of the target mRNA by blocking initiation or ribosome translocation.

We provide evidence that a significant portion of cytoplasmic dFMRP colocalizes with AGO1 but not GW182 (Fig. 3). This relationship is supported by co-immunoprecipitation data which suggests that dFMRP-containing complexes contain higher levels of AGO1 than GW182 (Fig. 3D). Based on these data, we propose that most dFMRP is interacting with a pool of AGO1 that is not loaded with miRNAs. GW182 proteins have an increased affinity for AGOs after miRNA loading (Elkayam et al. 2017). This suggests that GW182 is recruited to the miRISC once the AGO proteins are primed for target mRNA recognition. FMRP has also been shown to reversibly interact with Dicer-containing complexes (Cheever and Ceman 2009). After processing pre-miRNAs into mature miRNAs, Dicer transiently associates with AGOs while one strand is preferentially loaded (Miyoshi et al. 2009). Together, our data suggests that dFMRP is associated with AGO1 before (predominantly), during, and after biogenesis of the functional miRISC.

GW182 is an evolutionarily conserved core component of the miRISC in metazoans. It has been shown to interact directly with the AGO proteins and acts as a scaffold to aid in the recruitment of the CCR4-NOT deadenylase complex and, subsequently, the decapping enzymes (Niaz and Hussain 2018). Deadenylation followed by decapping triggers 5’-to-3’ mRNA decay. Interestingly, our data suggests that dFMRP, AGO1, and GW182 can all repress translation of a reporter mRNA that is not able to be deadenylated (Fig. 1E) (Kuzuoglu-Ozturk et al. 2016). Moreover, dFMRP represses reporter translation with no changes in mRNA levels (Fig. 1C). Taken together, these data suggest that dFMRP, AGO1, and GW182 repress reporter translation via a deadenylation- and decay-independent mechanism. GW182 has also been shown to block translation, in the absence of mRNA decay, by interacting with poly(A)-binding proteins (PABPs) (Niaz and Hussain 2018). Mechanistically, the GW182/PABP interaction is thought to interfere with mRNA circularization and thereby reduce the efficiency of translational initiation (Fabian et al. 2009; Zekri et al. 2009). Alternatively, the GW182/PABP interaction may reduce the affinity of PABP for the poly(A) tail. While we cannot rule out the idea that a FMRP/AGO1/GW182 complex is inhibiting initiation, our data suggests that it may instead be blocking elongation (Fig. 1D).

Our data also shows that dFMRP is capable of binding directly to the short BoxB stem loop (Fig. 4). Binding of FMRP to stem loop structures in target RNAs is not unprecedented. First, human FMRP can bind directly to a short stem loop in the noncoding Brain Cytoplasmic 1 (*BC1*) RNA (Zalfa et al. 2005). Second, FMRP can bind directly to a three tandem stem loops in the mRNA encoding for mouse Superoxide Dismutatase 1 (Sod1) and promote its translational activation (Bechara et al. 2009). Finally, mammalian FMRP has been shown to bind to more complex secondary structures in target RNAs including G-quadraplexes through its RGG box and RNA pseudoknot structures (kissing complexes) through its KH2 domain (Darnell et al. 2001; Schaeffer et al. 2001; Darnell et al. 2005). Interestingly, while G-quadraplexes are the best characterized structural element bound by mammalian FMRP, data suggests that *Drosophila* FMRP does not bind with high affinity to the strong G-quadraplex structure found in the *sc1* RNA (Darnell et al. 2009). This study also determined that the RGG box in dFMRP is poorly conserved relative to its mammalian orthologs. Collectively, these data suggest that dFMRP may not bind to target RNAs through an RGG/G-quadraplex interaction. It is possible that the RGG box in dFMRP may have deviated in its binding affinity, perhaps for small stem loop structures. Future work is needed to investigate if this mechanism is involved in the regulation of endogenous mRNAs.

## MATERIALS AND METHODS

### DNA constructs

For luciferase assays, the 5xBoxB FLuc reporters, RLuc plasmid, plasmids expressing λN-HA tagged proteins (dFMRP, AGO1, and GW182), and miR-958 have all be described previously (Eulalio et al. 2008; Nesler et al. 2013; Zekri et al. 2013; Starke et al. 2022). To construct the 3x, and 1x plasmids, we removed the 5XBoxB sequence from the pAc5.1C-FLuc-Stop-5BoxB plasmid (Addgene #21301) between the EcoRI and XhoI sites and replaced with oligos encoding for the 3xBoxB or 1xBoxB stem loop sequences. The 0xBoxB plasmid was generated by blunting the EcoRI and XhoI sites and then recircularizing the plasmid. For S2 cell colocalization experiments, GFP-tagged dFMRP has been previously described (Starke et al. 2022). The single isoform of DCR1 was amplified by RT-PCR from mRNA isolated from S2 cells and reverse transcribed using the Oligo(dT)-primed RNA to cDNA EcoDry premix (Takara Bio). The CDS for AGO1 and GW182 were PCR amplified from pAFW-Ago1 (Addgene #50553) and LD47780 (DGRC) respectively. All three were cloned downstream of mCherry in pAc5.1A (Invitrogen). For the co-immunoprecipitation experiments, the pAc5.1-λN-HA:dFMRP and pAFW-AGO1 plasmids have been previously described (Kawamata et al. 2009; Starke et al. 2022). To construct the FLAG-tagged GW182 plasmid, the CDS for GW182 was PCR amplified from LD47780 (DGRC) and inserted into the pAFW vector (DGRC) by Gateway Cloning (Invitrogen). For the EMSA experiments, the dFMRP CDS was PCR amplified from pAc5.1-EGFP-dFMRP and transferred into pET-His6-MBP-TEV (Addgene #29656) by ligation-independent cloning following QB3 Macrolab protocols (https://qb3.berkeley.edu/facility/qb3-macrolab/). DNA sequences for all oligonucleotides used for PCR throughout this study are provided in Supplementary Table 1.

### Cell culture and luciferase assays

*Drosophila* S2 cells (S2-DRSC; DGRC #181) were maintained at 25°C in 75 cm^2^ cell culture-treated flasks in M3 media (Sigma) supplemented with 10% FBS (Gibco). Transfections were carried out in 12-well plates using Effectene transfection reagent (Qiagen). Unless otherwise indicated (for gradient experiments), the transfection mixture (per well) contained 0.05 μg of the FLuc reporter plasmid, 0.2 μg of the RLuc transfection control plasmid, and 0.5 μg of the plasmid expressing the λN-HA control, λN-HA-tagged, or untagged proteins. For RNAi experiments, PCR primers containing a 5’ binding site for T7 RNA polymerase were designed using SnapDragon (https://www.flyrnai.org/snapdragon) and used to amplify sequences targeting each gene. dsRNA was synthesized from PCR product using a MEGAscript T7 kit (Ambion). 1 μg of dsRNA (per well) was added to each transfection mixture. All transfections were done in triplicate and luciferase activity measured after 3 days using the Dual Luciferase Reporter Assay System (Promega).

### Western blotting and quantitative real time PCR (qPCR)

Western blotting was done essentially as previously described (Starke et al. 2022). Antibodies targeting dFMRP (dilution 1:1000; Abcam #10299), DCR1 (dilution 1:1000; Abcam #4735), AGO1 (dilution 1:2000; Abcam #5070), GW182 (dilution 1:2000) (Schneider et al. 2006), HA (dilution 1:500; Cell Signaling Technology #2367), FLAG (dilution 1:1000; Sigma M2), and β-actin (dilution 1:1000; Abcam #8224). Bound primary antibodies were detected with horseradish peroxidase-coupled secondary antibodies (Cell Signaling Technologies) and then visualized by chemiluminescence (using SuperSignal West Chemiluminescence kits from Thermo Fisher).

Quantification of RNA abundance in S2 cells was done by qPCR. On the day of luciferase assays, total RNA was isolated from the remaining cells using TRIzol (Invitrogen) followed by column purification using the RNeasy Mini Kit (Qiagen) followed by DNAse treatment. 1 μg of total RNA was reverse transcribed using the double primed RNA to cDNA EcoDry premix (Takara Bio). Primers were designed targeting FLuc, Rluc, and β-actin as a control and analyzed on an IQ5 Thermal Cycler (BioRad) using the iQ SYBR Green Supermix (BioRad). All reactions were done in triplicate. To evaluate the specificity of PCR amplification for each primer set, we performed melt curve analysis. For relative quantification of transcript levels, we used the ΔΔCt method.

### Polysome profiling

S2 cells were transfected as described for luciferase assays except plasmid concentrations were doubled for 6-well plates. Three wells were transfected for each set of plasmids. After 3 days, cells were incubated with cycloheximide at a final concentration of 100 μg/ml for 10 min at 25°C. Cells were then pooled (∼ 10 x 10^6^ cells total), washed in ice cold PBS with cycloheximide, resuspended in polysome lysis buffer (10 mM HEPES (pH 7.4), 10 mM MgCl_2_, 150 mM KCl, 0.5% NP-40, 0.5 mM DTT, 100 U/ml SUPER-RNAse inhibitor (Thermo Fisher), complete EDTA-free protease inhibitor (Roche), and 100 μg/ml cycloheximide), and then lysed using a dounce homogenizer. Cell debris was removed by centrifugation at 14,000*g* at 4°C. 300 μl of each supernatant was layered on top of a linear 10-50% sucrose gradient in 10 mM HEPES 10 mM HEPES (pH 7.4), 15 mM MgCl_2_, 150 mM KCl). Centrifugation was done using an SV-41 rotor (Beckman) for 2h 30 min at 37,000 rpm at 4°C. Polysome profiles were measured by absorbance and fractions collected using a Gradient Station and Fractionator (Biocomp). For RNA analysis, 200 μl of each fraction was used. RNA was purified using TRIZOL followed by column purification using the DirectZol RNA mini kit (Zymo Research). Fractions were pooled (proportionally) based on polysome profiles into RNP, 40S ribosome, 60S ribosome, monosome, and polysomes (Supplemental Fig. S1). RNA was then reverse transcribed using the double primed RNA to cDNA EcoDry premix (Takara Bio). RNA abundance was determined by qPCR using primers targeting FLuc. Fold enrichment was calculated relative to the RNP sample. For protein analysis, protein was precipitated from 200 μl of each fraction with TCA and pellets solubilized in 100 μl of 2x Laemmli sample buffer for 10 min at 95°C. 15 μl of each sample was loaded onto a 4-20% Mini Protean gel (BioRad), separated, and analyzed by Western blot.

### Co-immunoprecipitation

S2 cell transfections were carried out in 6-well plates using Effectene transfection reagent (Qiagen). Each well was transfected with 1 μg of the pAc5.1-λN-HA:dFMRP and 1 μg pAFW-AGO1 or pAFW-GW182 plasmids. After incubation at 25°C for 3 days, cells were resuspended in ice cold Pierce IP Lysis Buffer (25 mM Tris (pH = 7.4), 150 mM NaCl, 1mM EDTA, 1% NP-40, and 5% glycerol). Cell debris was removed by centrifugation at 13,000*g* for 10 min at 4°C. HA-tagged FMRP (and associated proteins) was immunoprecipitated using the HA-Tag IP/Co-IP kit (Pierce). Protein was dissociated from magnetic beads by incubation with 50 μl of 2x Laemmli sample buffer for 10 min at 95°C and subjected to SDS-PAGE followed by Western analysis.

### S2 cell imaging and colocalization

S2 cell transfections were carried out in 6-well plates using Effectene transfection reagent (Qiagen). Each well was transfected with 1 μg of the pAc5.1-EGFP:dFMRP and 1 μg of pAc5.1-mCherry plasmids containing DCR1, AGO1, or GW182. After incubation at 25°C for 3 days, cells were resuspended and 200 μl transferred to poly-D-lysine coated #1.0 glass bottom dishes (Cellvis) and allowed to settle for ∼ 5 min. Live cell fluorescence images were obtained using an Olympus FV3000 scanning confocal microscope with a 100x (N.A. = 1.4) objective digitally zoomed to 2.95 (the optimal setting for this objective per the Fluoview software). To determine the degree of colocalization, single focal planes were obtained for 12 to 14 cells and analyzed in FIJI/ImageJ2 using the JACoP plugin (Bolte and Cordelieres 2006). Images were cropped to the smallest area possible to eliminate colocalization outside of the cell of interest. In JACoP, Pearson’s and Mander’s coefficient results were generated and recorded for statistical analysis.

### Protein purification and electrophoretic mobility shift assay

BL21-DE3 cells (New England Biolabs) were transformed with pET-His6-MBP-TEV-dFMRP and protein expression induced by incubation with IPTG. After ∼ 4 hours of expression, cells were lysed in lysis buffer (50 mM Tris (pH = 8), 100 mM NaCl, 5 mM β-mercaptoethanol, 5% glycerol, and 1mM imidazole). MBP-tagged protein was purified by affinity column chromatography using Ni-IMAC resin (Thermo Fisher) followed by size exclusion chromatography using a Superdex 200 column (Sigma Aldrich). Protein was eluted off columns and then analyzed SDS-PAGE followed by Coomassie staining (Supplemental Figure S3). The remaining protein was run through a 50 kDA cut off spin concentrator (Millipore) and dialyzed into 2X storage buffer (48 mM HEPES, 500 mM NaCl, 20% glycerol, and 4 mM DTT). The sample was further diluted by half with molecular grade 100% glycerol (Sigma Aldrich) and stored at −80°C prior to use.

For EMSA experiments, short RNAs corresponding to the 17 nucleotide BoxB stem loop or the A14 unstructured control were synthesized *in vitro* and purified as we have previously described (Langeberg et al. 2020). Single stranded RNA and purified full-length FMRP were mixed and then allowed to bind at room temperature for 20 minutes. There was 1 µg of RNA added to each binding reactions for both the BoxB (MW = 5805 for 5’ OH) and A14 (MW = 4626 for 5’ OH). Molar ratios indicated are protein to RNA. After incubation, samples were separated by electrophoresis on a 20% native polyacrylamide gel. Following electrophoresis, these “cold” native gels were stained with methylene blue and RNA bands imaged for analysis.

### Fly stocks, NMJ dissections, immunofluorescence, and quantification

For all experiments, both male and female flies were used for analysis. All crosses were incubated at 25°C with 12-hour light/dark cycles and 60% humidity on standard Bloomington media. Flies used in this study were *w^1118^ (Iso31), dFmr1^Δ113^*, *Ago1^K00208^* (Bloomington *Drosophila* Stock Center) and *gw^1^* (Schneider et al. 2006). NMJ dissections were done essentially as previously described (Patel et al. 2020). Larval body wall preps were prepared by dissecting wandering 3^rd^ instar larvae in Ca^2+^-free HL3 saline. For imaging the NMJ, larval preps were fixed with 3.5% paraformaldehyde in PBS and then immunostained with antibodies targeting presynaptic horseradish peroxidase (HRP) and postsynaptic discs large (DLG). The specific antibodies used were Alexa 568-conjugated anti-HRP (1:1000, Jackson Immunoresearch), mouse anti-DLG (1:100, Developmental Studies Hybridoma Bank), and an anti-mouse Alexa 488-conjugaged secondary (1:1000, Molecular Probes). Preps were then mounted on charged slides in DAPI Flourmount G (Southern Biotech) and images were obtained using an Olympus FV3000 scanning confocal microscope with a 60x (N.A. = 1.42) objective. The number of type 1b synaptic boutons at muscles 6 and 7 (m6/7) in abdominal segment 3 (A3) were manually quantified using the built in cell counting plugin in FIJI/ImageJ2 as previously described (Pradhan et al. 2012). Boutons were defined as a distinctive swelling at the NMJ marked by both DLG and HRP and distinguished from type 1s boutons based on size and a greater amount of DLG staining. To account for differences between genotypes in the scaling of NMJs to muscle size, synaptic bouton numbers were normalized to muscle surface area (MSA). MSA was calculated from images of the corresponding m6/7 obtained with a 20x objective (N.A. = 0.85) and quantified in FIJI/ImageJ2. Data was collected from 18 larvae for each of the indicated genotypes for statistical analysis.

## SUPPLMENTAL MATERIAL

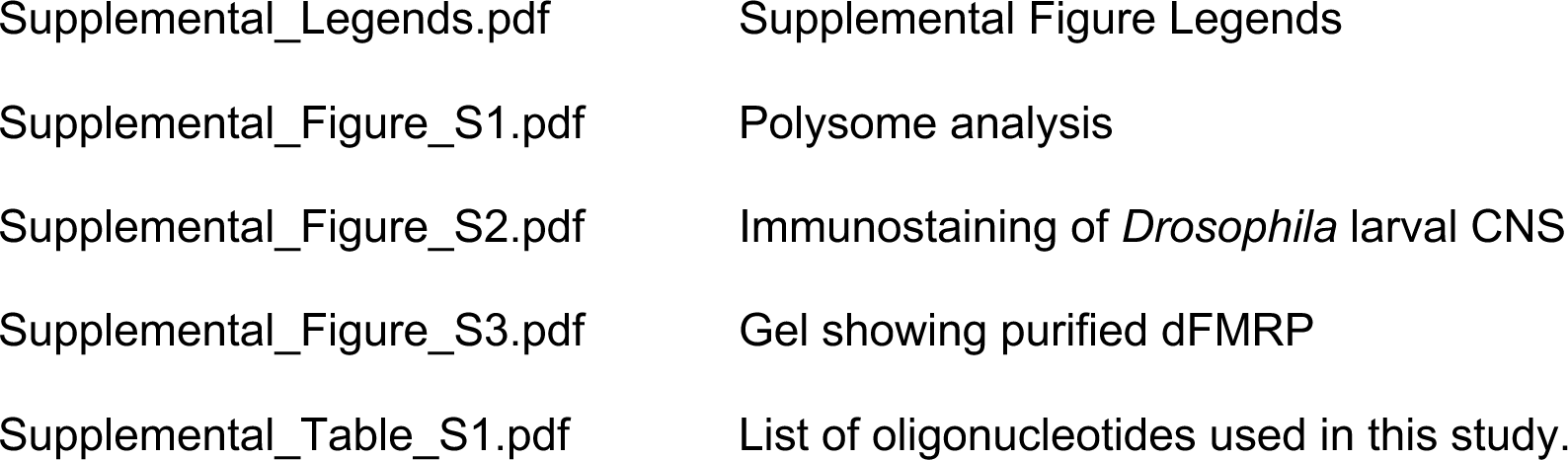

## ACKNOWLEDGEMENTS

We would like to thank members of the Barbee lab for useful discussions. The deadenylation- and scanning-independent reporter plasmids were a kind give from Dr. Elisa Izaurralde. Plasmids not developed for use in this study were obtained from Addgene and the *Drosophila* Genomics Resource Center. S2 cells were purchased from the *Drosophila* Genomics Resource Center. The *dFmr1* and *Ago1* fly lines were obtained from the Bloomington *Drosophila* Stock Center. The *GW182* fly line and antibody was a generous gift from Andrew Simmonds.

