## Supplemental Data for "*Drosophila* FMRP recruits the miRISC to target mRNAs to repress translation"

**SUPPLEMENTAL FIGURE LEGENDS:**

**Supplemental Figure 1. Polysome analysis.** (A-B) Graphs showing polysome profiles and fractions collected (including those pooled for qPCR analysis in Fig. 1) for S2 cells transfected with the FLuc-5xBoxB reporter and either the λN-HA peptide (A) or λN-HA-tagged dFMRP (B).

**Supplemental Figure 2. Colocalization of dFMRP with AGO1 and GW182 in the larval CNS.** Representative images of third instar larval ventral ganglia immunostained with antibodies targeting dFMRP (green) counterstained with antibodies targeting AGO1 or GW182 (red). Merged images show a significant degree of coexpression in the cell body of larval neurons (yellow).

**Supplemental Figure 3. Purified His6-MBP-tagged dFMRP.** Image of a Coomassie-stained gel showing dFMRP protein after purification by affinity column chromatography followed by size exclusion chromatography. The predicted size of the isoform used in this study (*dFmr1-PD*) is ~ 75kDa and is indicated by the arrow. Note that addition of the TEV protease results in the internal cleavage and fragmentation of most full-length dFMRP. Likely cleavage products are indicated by an asterisk. Only full length dFMRP was used for the EMSA experiments shown in Fig. 4.

Figure S1

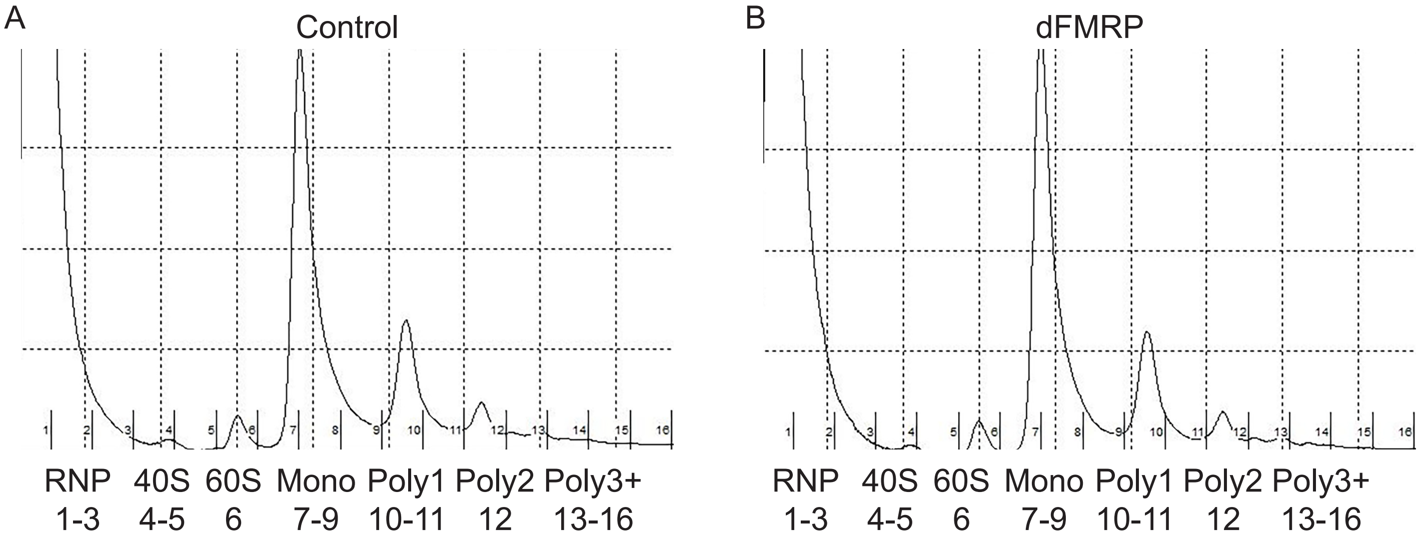
A
Control
RNP 1-3
40S 4-5
60S 6
Mono 7-9
Poly1 10-11
Poly2 12
Poly3+ 13-16
B
dFMRP
RNP 1-3
40S 4-5
60S 6
Mono 7-9
Poly1 10-11
Poly2 12
Poly3+ 13-16

Figure S2

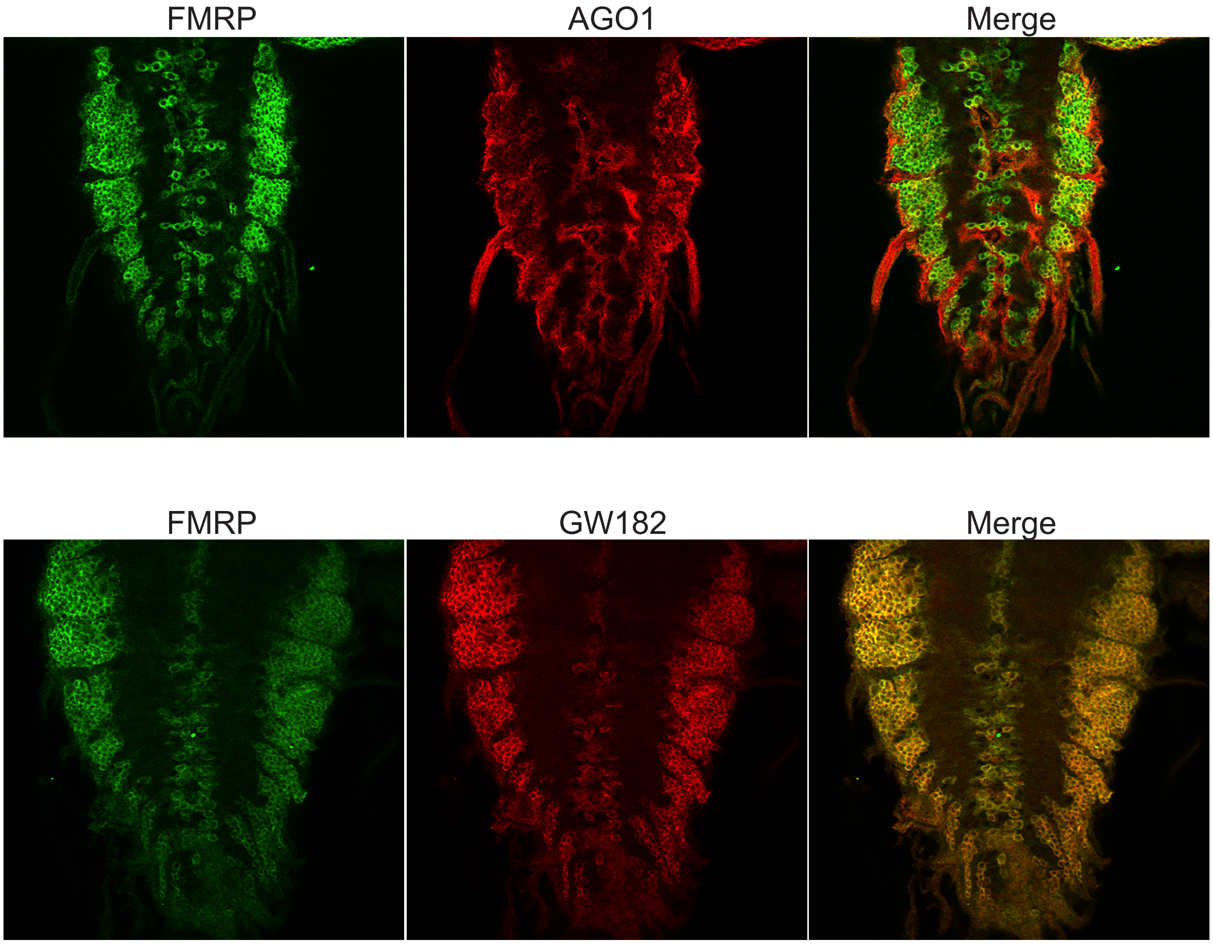

Figure S3

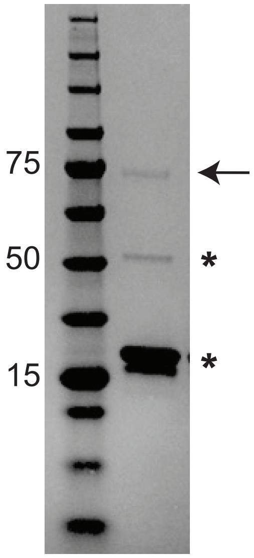

| **Table S1. List of Oligonucleotides used in this study** | |
|---|---|
| **Oligonucleotide** | **Sequence (5' to 3')** |
| 3xBoxB cloning oligo | GAATTCGGGCCCTGAAGAAGGGCCCATATAGGGCCCTGAAGAAG GGCCCATATAGGGCCCTGAAGAAGGGCCCCTCGAG |
| 1xBoxB cloning oligo | GAATTCGGGCCCTGAAGAAGGGCCCCTCGAG |
| Dicer1 cloning forward primer | GGTGGTGCGGCCGCATGGCGTTCCACTGGTGCGAC |
| Dicer1 cloning reverse primer | GGTGGTCCGCGGTTAGTCTTTTTTGGCTATCAAGCC |
| AGO1 cloning forward primer | ATGTCCACGGAGCGTGAG |
| AGO1 cloning reverse primer | TTAGGCAAAGTACATGACCTTCTTG |
| GW182 cloning forward primer | ATGGCGTTCCACTGGTGC |
| GW812 cloning reverse primer | TTAGTCTTTTTTGGCTATCAAGCCC |
| FMRP Lic cloning forward primer | TACTTCCAATCCAATGCAGAAGATCTCCTCGTGGAAGTTCGGC |
| FMRP Lic cloning reverse primer | TTATCCACTTCCAATGTTATTAGGACGTGCCATTGACCAGGCC |
| *Dcr1* RNAi forward primer | TAATACGACTCACTATAGGGCGGAACACGATTATTTGCCT |
| *Dcr1* RNAi reverse primer | TAATACGACTCACTATAGGGCGCAACACGGTGACAATATC |
| *Ago1* RNAi forward primer | TAATACGACTCACTATAGGGATTTGATTTCTATCT ATGCAGCCA |
| *Ago1* RNAi reverse primer | TAATACGACTCACTATAGGGGCCCTGGCCATGGCACCTGGCGTA |
| *GW182* RNAi forward primer | TAATACGACTCACTATAGGGAATCCAAGTAATCCTATAAGCAG |
| *GW182* RNAi reverse primer | TAATACGACTCACTATAGGGATTGCTTGCTTTGCTTAATGA |
| EGFP RNAi forward primer | TAATACGACTCACTATAGGGAAGAAATCCTCGCCCATTAC |
| EGFP RNAi reverse primer | TAATACGACTCACTATAGGGTGCTCAGGTAGTGGTTGTCG |
| FLuc reporter qPCR forward primer | AAACGCTTCCACCTACCAGG |
| FLuc reporter qPCR reverse primer | TGATCAGAATGGCGCTGGTT |
| RLuc reporter qPCR forward primer | TTACATGGTAACGCGGCCT |
| RLuc reporter qPCR forward primer | TAATACACCGCGCTACTGGC |
